# Alpha Activity Reflects the Magnitude of an Individual Bias in Human Perception

**DOI:** 10.1101/759159

**Authors:** Laetitia Grabot, Christoph Kayser

## Abstract

Biases in sensory perception can arise from both experimental manipulations and personal trait-like features. These idiosyncratic biases and their neural underpinnings are often overlooked in studies on the physiology underlying perception. A potential candidate mechanism reflecting such idiosyncratic biases could be spontaneous alpha band activity, a prominent brain rhythm known to influence perceptual reports in general. Using a temporal order judgement task, we here tested the hypothesis that alpha power reflects the overcoming of an idiosyncratic bias. Importantly, to understand the interplay between idiosyncratic biases and contextual (temporary) biases induced by experimental manipulations, we quantified this relation before and after temporal recalibration. Using EEG recordings in human participants (male and female), we find that pre-stimulus frontal alpha power correlates with the tendency to respond relative to an own idiosyncratic bias, with stronger alpha leading to responses matching the bias. In contrast, alpha power does not predict response correctness. These results also hold after temporal recalibration and are specific to the alpha band, suggesting that alpha band activity reflects, directly or indirectly, processes that help to overcome an individual’s momentary bias in perception. We propose that combined with established roles of parietal alpha in the encoding of sensory information frontal alpha reflects complementary mechanisms influencing perceptual decisions.

**Significance statement:** The brain is a biased organ, frequently generating systematically distorted percepts of the world, leading each of us to evolve in our own subjective reality. However, such biases are often overlooked or considered noise when studying the neural mechanisms underlying perception. We show that spontaneous alpha band activity predicts the degree of biasedness of human choices in a time perception task, suggesting that alpha activity indexes processes needed to overcome an individual’s idiosyncratic bias. This result provides a window onto the neural underpinnings of subjective perception, and offers the possibility to quantify or manipulate such priors in future studies.

## Introduction

Perception is not only driven by the incoming sensory information but is also shaped by expectations, knowledge and other individual biases. These can arise from both temporary contextual effects and long-term priors. Temporary biases can be experimentally manipulated by changing the probability of one response over another, or through contextual recalibration (Summerfield & Egner, 2009; de Lange, Heilbron, & Kok, 2018). The long-term priors comprise life-long learned and stable assumptions about the world, such as the sun shining from above (Sun & Perona, 1998). These priors may vary between individuals, but are stable for a given individual, and reflect idiosyncratic biases. These are often unknown to the experimenter and may be shadowed by experimental manipulations or regarded as inter-individual noise eliminated during data analysis (Kanai & Rees, 2011; Wexler et al., 2015; Grabot & Wassenhove, 2017; Rahnev & Denison, 2018; Lebovich et al., 2019). We here argue that such inter-individual variability in temporary and idiosyncratic biases provides key perspectives on the neural mechanisms underlying perception. Particularly, we ask whether pre-stimulus oscillatory activity reflects the neural underpinnings of how such idiosyncratic biases affect perception, and if so, how these interact with those reflecting temporary biases.

The influence of prior expectations and knowledge on perceptual decisions is often thought to arise from high-level prefrontal and parietal regions, and is conveyed to sensory areas via top-down mechanism (Engel, Fries, & Singer, 2001; Summerfield & Lange, 2014; de Lange, Heilbron, & Kok, 2018). One neural signature supposedly reflecting these underlying processes is alpha band activity (van Kerkoerle, et al., 2014; Michalareas, et al., 2016; Sherman, Kanai, Seth, & VanRullen, 2016; Mayer, Schwiedrzik, Wibral, Singer, & Melloni, 2016), generally known to be predictive of upcoming perceptual decisions (Ergenoglu, et al., 2004; Hanslmayr, et al., 2007; van Dijk, Schoffelen, Oostenveld, & Jensen, 2008; Mathewson, Gratton, Fabiani, Beck, & Ro, 2009). Recent studies have suggested that pre-stimulus alpha activity may reflect the criterion used to commit a specific response and may hence reflect a perceptual or decisional bias (Limbach & Corballis, 2016; Iemi, Chaumon, Crouzet, & Busch, 2017; Craddock, Poliakoff, El-deredy, Klepousniotou, & Lloyd, 2017; Iemi & Busch, 2018; Rohe, Ehlis, & Noppeney, 2019). Along such a role in perceptual decision-making, alpha activity was shown to correlate with subjective awareness (Benwell, et al., 2017; Lange, Oostenveld, & Fries, 2013; Gulbinaite, İlhan, & VanRullen, 2017) and decision confidence (Samaha, Iemi, & Postle, 2017; Wöstmann, Waschke, & Obleser, 2018). Still, it remains unclear whether pre-stimulus activity indeed reflects processes shaping an individual’s intrinsic bias or processes facilitating veridical sensory encoding, as previous work did not unambiguously quantify the relation of spontaneous brain activity to idiosyncratic and temporary biases.

We have previously proposed the hypothesis that alpha band activity compensates for an individual’s idiosyncratic bias: thereby low alpha power enhances the probability to respond against the bias, while high power increases the tendency to follow the bias (Grabot & Wassenhove, 2017; Grabot, Kösem, Azizi, & Wassenhove, 2017). To directly test this hypothesis, we here used a temporal order judgement task, in which perception is shaped by idiosyncratic biases (Freeman, et al., 2013; Ipser, Karlinski, & Freeman, 2018; Grabot & Wassenhove, 2017) and can be manipulated by inducing temporary biases through temporal recalibration. Importantly, our paradigm was specifically designed to dissociate intrinsic biases from overall task performance. This allowed us to test whether pre-stimulus activity is predictive of the correctness or the degree of biasedness of a response, factors that were confounded in previous work. Our results confirm that a decrease in pre-stimulus alpha power enhances the probability to respond against the individual idiosyncratic bias and that inter-individual fluctuations in alpha power correlate with the magnitude of this perceptual bias. Furthermore, by testing the influence of contextual recalibration (Fujisaki, Shimojo, Kashino, & Nishida, 2004; Van der Burg, Alais, & Cass, 2013) we show that this relation persists after temporal recalibration, suggesting that alpha power reflects both idiosyncratic and temporary biases.

## Materials & Methods

### Participants

Forty right-handed naive participants with normal or corrected-to-normal vision and normal hearing were tested in a first screening session (10 male, 30 female; mean ± SD age = 24 ± 3 years). Each provided written informed consent in accordance with the Declaration of Helsinki (World Medical Association, 2013), and the study was approved by the local ethics committee of Bielefeld University. The screening session was designed to collect a planned sample of 24 participants, balanced according to their preferred temporal order (see below), and with a negative JND1 and a positive JND2 (Just Noticeable Difference, defined as in Fig. 1B). 12 participants were excluded based on the screening data, including 10 whose JNDs fell outside the range of tested stimulus onset asynchronies (according to the criteria used in Spence et al., 2001), one whose JNDs were both negative, and one who did not complete the entire session. In total, 28 participants joined the EEG session, 4 of which had to be excluded because they had less than 20 trials in at least one condition. Thus, in the final analysis, we included 24 participants (7 male, 17 female; age = 24 ± 2 years mean ± SD).

**Fig. 1:**
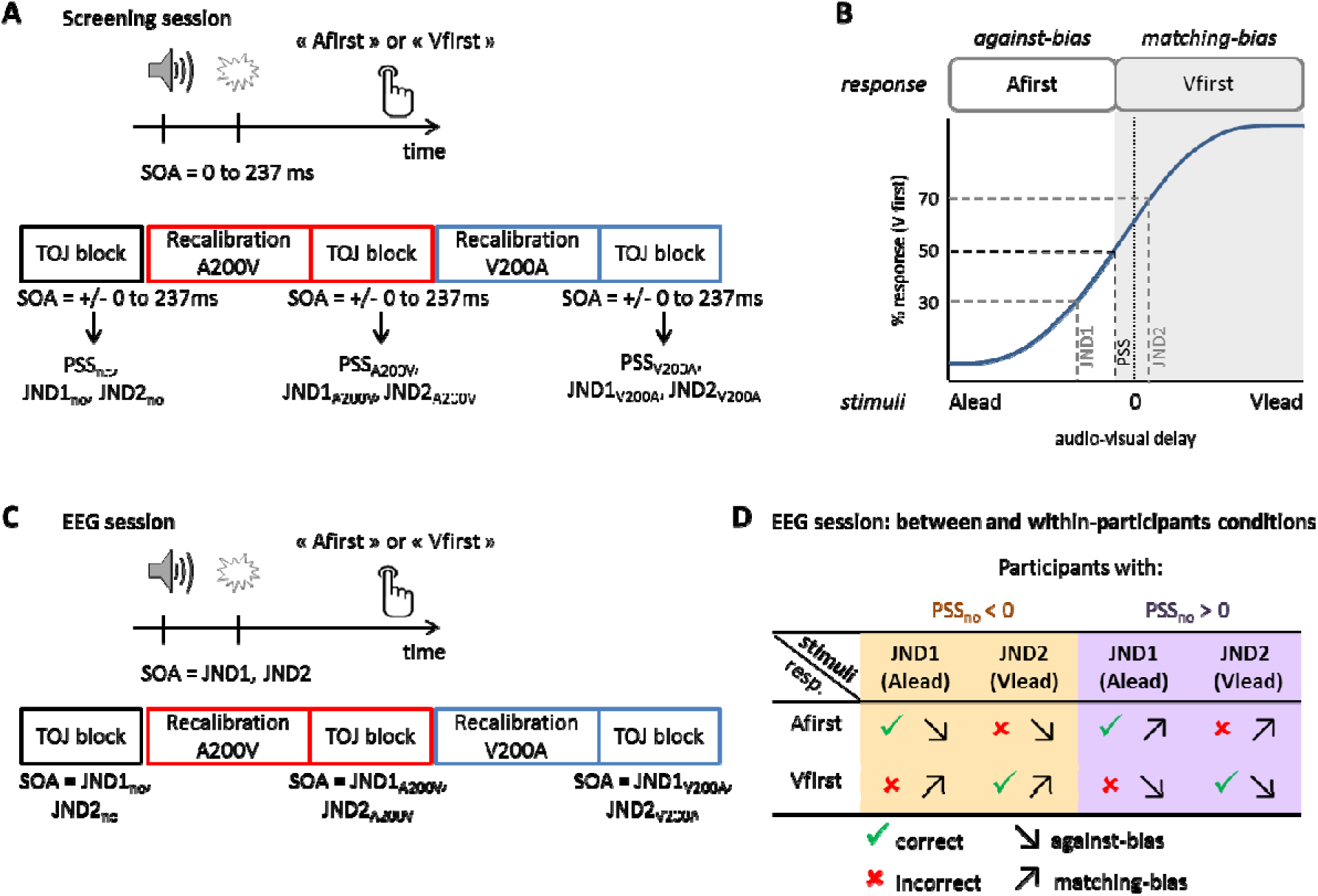
Experimental design. **A.** The screening session was based on a temporal order judgment task and comprised three conditions: no adaptation, audio-leading adaptation (A200V) and visual-leading adaptation (V200A). We assessed the individual bias in temporal order perception (PSS_no_), adapted PSSs (PSS_A200V_ and PSS_V200A_) and JNDs. **B.** Schematics of a psychometric curve obtained after the screening session. The PSS was defined as the SOA for which a participant’s performance was at chance level (50%). The JNDs were defined as the SOAs for which a participant’s performance was at 30 % (JND1) and 70% (JND2) of performance. If a participant has a negative PSS as shown here, i.e. the sound needs to be presented first so that s/he can perceive simultaneity, s/he has an increased tendency to reply Vfirst (matching-bias response) compared to Afirst (against-bias response). **C.** In the EEG session, the two JNDs derived per condition in the screening session were used as SOAs. **D.** The correctness and biasedness contrasts are fully orthogonal, and they comprise the same number of audio-leading and visual-leading pairs.

### Stimuli

The experiment was controlled using the Psychophysics Toolbox (Version 3.0.14; http://psychtoolbox.org/) and run using MATLAB (Version R2017a; The MathWorks, Inc., Natick, MA). The visual stimuli were white rings (outer diameter = 12°, inner diameter = 8°, 275 cd/m^2^) presented for 25 ms (three frames) and centered on a grey LCD screen (refresh rate = 120 Hz, resolution = 1920 × 1080 pixels, 16 cd/m^2^). Participants were asked to fixate a black cross presented in the middle of the screen during the whole experiment. Auditory stimuli were sine-wave tones (2 kHz) presented for 25 ms (including 5-ms fade-in and 5-ms fade-out) at a comfortable hearing level (73 dB). They were presented trough speakers placed behind each side of the monitor screen to ensure a perceived co-localization of sound and visual stimuli. The relative timing of auditory and visual stimuli was tested to be precise up to ± 3ms using an oscilloscope and a photodiode.

### Screening session

The experiment comprised a screening session and two EEG sessions based on a temporal order judgment (TOJ) task (Fig. 1A). The three sessions took place within approximately 10 days (7 ± 6 days between the screening session and the first EEG session, and 3 ± 2 days between the two EEG sessions). Experiments took place in a darkened and electrically shielded room (Desone Ebox; Germany), where the participant sat at 90 cm distance from the screen.

The screening session aimed at estimating the temporal thresholds associated with judging the temporal order of a pair of audiovisual stimuli (Fig. 1A, 1B). The individual thresholds were then used as stimulus onset asynchrony (SOA) in the following EEG sessions, ensuring a similar ratio of Afirst and Vfirst responses. In the screening session, the audiovisual stimuli were separated by 12 different SOAs: ± 20, ± 50, ± 80, ± 110, ± 160, ± 250 ms. Negative values indicate that the sound was presented first.

The task was performed in three conditions: without recalibration (no), after sound-leading recalibration (A200V) and after visual-leading recalibration (V200A). In the no recalibration condition, participants were presented with a pair of audiovisual stimuli and had to judge which of the two came first, in a two-alternative forced choice, by pressing a key on a keyboard. The trials in A200V- and V200A-recalibration conditions were split into 3 blocks per condition. Each block consisted of an exposure phase and a test phase. During the exposure phase, 84 audiovisual pairs with an identical SOA of 200 ms were presented at roughly 1 Hz (uniform distribution: 900-1100ms). In A200V-recalibration (resp. V200A) blocks, 80 trials were audio-leading (resp. visual-leading) while 4 deviant trials were visual-leading (resp. audio-leading). To enhance recalibration effects and to ensure that participants kept their attention focused they were asked to detect these deviant trials as fast as possible (Heron, Roach, Whitaker, & Hanson, 2010). The hit rate was on average 84 ± 10% and the false alarm rate was 1 ± 1 % (mean ± SD). The exposure phase was immediately followed by a test phase. Top-up trials were used to maintain recalibration (Fujisaki, Shimojo, Kashino, & Nishida, 2004; Vroomen, Keetels, Gelder, & Bertelson, 2004). We used a minimum number of top-up trials sufficient to enhance recalibration effects (Cai, Stetson, & Eagleman, 2012) and tested two consecutive audiovisual pair at once. Three top-up trials were presented at roughly 1Hz (uniform distribution: 900-1100ms). 700ms after the last top-up trial, the black fixation cross turned green, indicating to the participant to judge the order of the two next pairs of AV stimuli.

For all conditions (no, A200V, V200A), inter-trial interval delays were uniform between 900 and 1100 ms. In the screening session, each SOA was repeated 24 times per condition. The screening session consisted of no recalibration block, followed by three A200V recalibration and three V200A recalibration blocks, with randomized order. Participants were allowed to take a short break between blocks. Before the experiment, they were trained with 10 repetitions of maximal SOAs (± 250ms) for each recalibration condition and received feedback. Button mapping for responses was balanced between participants.

### PSS and JNDs estimation

In the screening session, we determined the order bias (PSS) and the just noticeable differences (JNDs) for each participant using the method of constant stimuli. For each recalibration condition (no, A200V and V200A), the percentage of Vfirst responses from the screening session was determined as a function of SOA and fit using a psychometric curve described by a logistic function with three parameters: bias, slope and lapse rate (Spence & Squire, 2003; Mégevand, Molholm, Nayak, & Foxe, 2013). We chose a three-parameter model based on a piloting test (n = 6) which revealed a lower Bayesian information and Akaike information criterion for this model compared to two- (bias and slope) or four-parameter (bias, slope, lapse rate for auditory-leading stimuli, lapse rate for visual-leading stimuli) models. Here, the goodness of fit (*R*^2^) was 90.2 ± 8.3% on average across participants (mean ± SD). The PSS, JND1 and JND2 were defined as the SOA for which a participant’s performance was respectively at chance level (50%), 30% and 70% (Fig. 1B) for each recalibration condition (no, A200V and V200A). To obtain a balanced design between correctness and bias (Fig. 1D), it was necessary that each individual’s JND1 was negative (sound-leading SOA), and each JND2 positive (visual-leading). To achieve this, we had to adjust the definition of JNDs for 5 participants (the JNDs performance ratios deviating from 30/70 for these participants were respectively: 23/70 in V200A condition; 30/78 in no condition, 30/87 in A200V condition, 30/78 in V200A condition; 30/93 in V200A condition; 30/95 in no condition, 30/82 in A200V condition, 30/90 in V200A condition; 30/92 in no condition, 30/75 in A200V condition).

### EEG sessions

Each EEG session consisted of two no recalibration, four A200V recalibration and four V200A recalibration blocks. Here, we used the three pairs of JND’s derived from the screening session as SOAs, and each SOA was repeated 200 times per condition (hence featuring two stimulus conditions per recalibration condition). The ITI preceding test trials was randomly drawn from a uniform distribution between 1.5-1.7s. Four catch trials (SOA = ± 250ms) were introduced in each block (∼4% of the total number of trials) to test the involvement of the participants. 88 ± 7 % (mean ± SD) of the catch trials were correctly detected. The no recalibration blocks were always presented first, followed by the 8 recalibration blocks in a randomized order.

### EEG acquisition and preprocessing

EEG signals were recorded using an active 128 channel Biosemi system (BioSemi, B. V., The Netherlands), with additional four electrodes placed near the outer canthi and below the eyes to record the electro-occulogram (EOG). Electrode offsets were below 25mV. Offline preprocessing and analysis were performed with MATLAB R2017a (The MathWorks, Natick, MA, USA) using the Fieldtrip toolbox (Oostenveld, Fries, Maris, & Schoffelen, 2011). The data were band-pass filtered between 0.2 and 90Hz, resampled to 50Hz and epoched from −1.5s before the first stimulus onset to 1s after. We removed ICA components that reflect eye movement artefacts, localized muscle activity or poor electrode contacts (12 ± 6 rejected components per block, mean ± SD). These were identified following definitions provided in the literature (Hipp & Siegel, 2013; O’Beirne & Patuzzi, 1999). Furthermore, trials with amplitude exceeding 150μV after 1Hz high-passed filtering and trials with reaction times shorter than 200ms were rejected. In total, 3 ± 5 % of trials (mean ± SD) were rejected across all participants. The EEG signals were not re-referenced to facilitate the analysis of local alpha power and phase effects (Yao, et al., 2005).

### Logistic modelling of behavioral data

To quantify the influence of the current stimulus and the previous stimulus or previous response on single-trial responses, we used generalized linear models using a binomial distribution with a logit link function. The model parameters were estimated using maximum likelihood methods based on Laplace approximation for each participant. The outcome variable was the response of the current trial (respN: Afirst=1, Vfirst=0), the predictors were the physical stimulus of the current trial (stimN: Alead=1, Vlead=-1), the physical stimulus of the previous trial (stimN-1: Alead=1, Vlead=-1) and the response of the previous trial (respN-1: Afirst=1, Vfirst=-1). All variables were treated as categorical variables. We first determined which behavioral model was explaining best the variance of the dataset by comparing Bayesian Information Criterion (BIC) after summing it across participants (Table 1).

**Table 1.** Models comparison for the trial-by-trial recalibration analysis. The table lists the group-averaged Bayesian Information Criteria (BIC) differences from the best model (model 3, red). The BIC were summed across participants.

| model | BIC |
| --- | --- |
| 1 | 1341 |
| 2 | 1115 |
| 3 | 0 |
| 4 | 73 |
| 5 | 39 |
| 6 | 112 |
| 7 | 202 |
| 8 | 210 |

### Time-frequency analysis

We extracted the single-trial power in different frequency bands between −1.5s to 0.5s around the first stimulus onset. To avoid any post-stimulus contamination, each trial was windowed (with a function equal to zero after stimulus onset and equal to 1 before, with a 27.5ms transition realized by a Hann window). The alpha power was obtained by using Morlet wavelets (10Hz, 3.5 cycles, which gives a spectral bandwidth of 7.1-12.8Hz and a temporal resolution of 350ms) and log-transformed. We ensured that the individual alpha peak frequency (IAF) matched the chosen generic alpha band, by extracting the global IAF from the power spectrum density computed on a −1.5 to −0.1s pre-stimulus time window. These ranged from 7.1 to 11.4Hz (average +/- SD = 9.8 +/- 1.3Hz, median = 10.0Hz). We also extracted the power from theta (6Hz, 3.5 cycles, spectral bandwidth = 4.3-7.7Hz), low beta (17Hz, 5 cycles, spectral bandwidth = 13.6-20.4Hz) and high beta bands (25Hz, 5 cycles, spectral bandwidth = 20-30Hz).

To investigate whether pre-stimulus alpha power was predictive of response correctness or biasedness, the alpha power was z-scored for each participant using the average and standard deviation across all trials and time points between −600 and −200ms for each sensor. We ignored the 200ms just prior to stimulus onset, as this time period is affected (given the wavelet length) by the windowing procedure, and we aimed to reduce any post-stimulus influence on spectral estimates as much as possible. The trials were then split into two groups depending on the contrast of interest (correct vs incorrect response, matching-bias vs against-bias responses). On average across participants, there were 551 ± 71 trials in the correct condition, 228 ± 70 in the incorrect, 397 ± 72 in the matching-bias condition and 382 ± 61 in the against-bias condition (mean ± SD). For the correlation analyses with the PSS, we used the raw alpha power differences between Afirst and Vfirst responses without baseline or z-scoring to be able to compare absolute changes in alpha between participants.

### Source reconstruction

We performed source reconstruction on the pre-stimulus alpha activity (−600 to −200ms) using DICS and re-referenced data (Gross, et al., 2001). Cross-spectral density matrices were computed using multiple tapers in the alpha band (centered in 10Hz, spectral smoothing of 3Hz). We created a forward model by using a standard 3D source model with 6mm dipole spacing and a MRI template from Fieldtrip, on which we manually aligned the 128-electrode array. We then computed the common inverse filter from all trials using DICS and applied this filter to each condition.

### Phase analysis

We used the phase opposition sum index (POS) to determine whether the instantaneous pre-stimulus alpha phase was concentrated around a mean value statistically different between two conditions (VanRullen, 2016). For this, we equalized the number of trials for conditions of each contrast, leading to 228 ± 68 trials for the correctness contrast and 341 ± 45 for the biasedness contrast (mean ± SD across participants). We computed POS index, based on the inter-trial coherence (ITC), for each condition, sensor and time point between −600 and - 200ms, as follows:

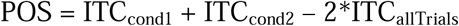

### Statistical analyses

The power difference (−600 to −200ms) between conditions was tested using a non-parametric cluster-based permutation procedure based on paired-sample t-tests (Fig. 2A) using the following parameters: two-sided t-test, alpha level for thresholding individual points at p=0.05, minimal number of neighbors in a cluster of 3, t-statistics performed on the maximal sum across cluster, 4000 randomizations. The same procedure was used to assess the significance of the Pearson correlation.

**Fig. 2.**
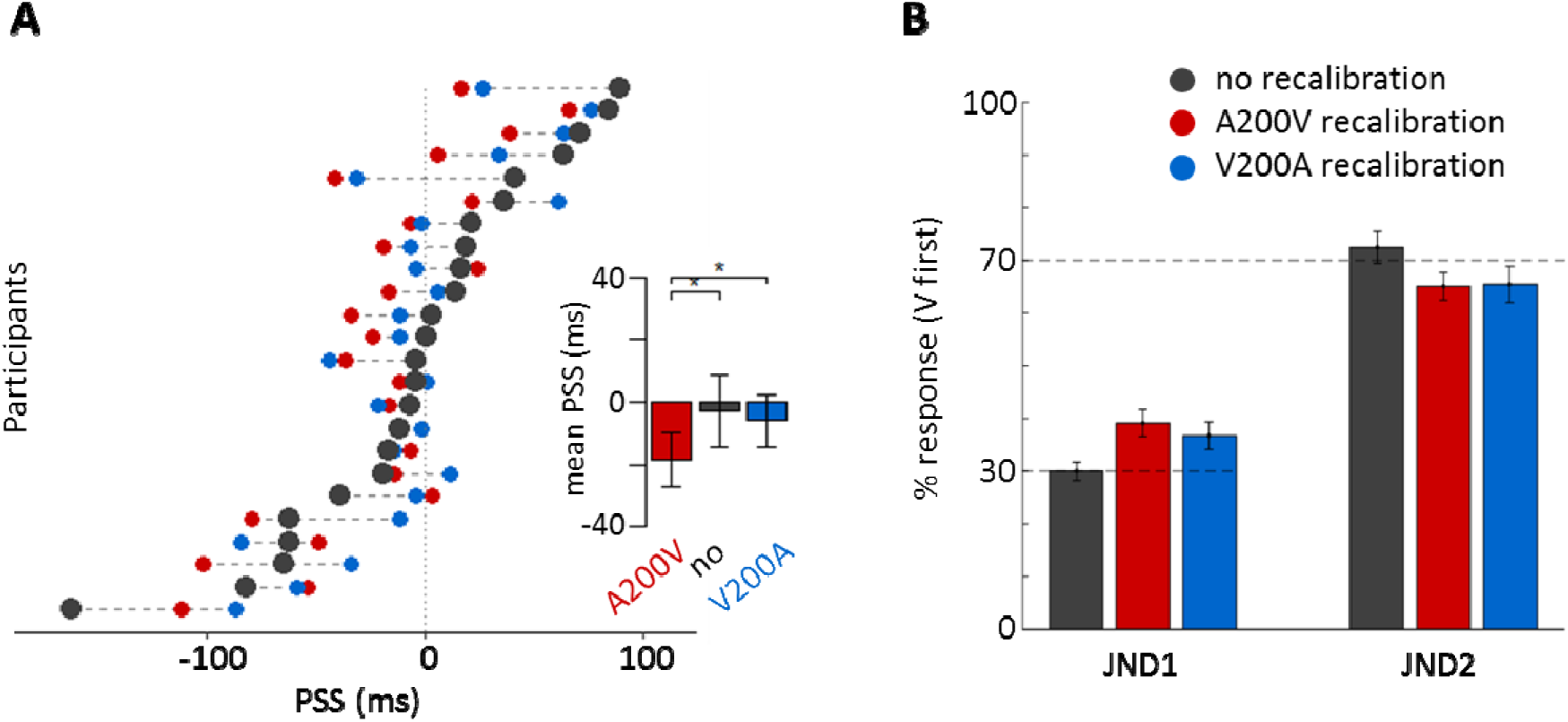
Behavioral results. **A.** Each line shows the participant’s PSSs for each condition (grey: no, red: A200V, blue: V200A). The insert depicts the group-averaged PSS for each condition (error bars shows ± 1SEM). Asterisks indicate the significance level of the repeated-measure one-way ANOVA (with *p < 0.05) **B.** Behavioral results of the EEG session. The percentage of flash-first responses is shown for each SOA (JND1 and JND2) and adaptation condition (error bars show ± 1SEM).

The phase difference between conditions was statistically assessed using a permutation-based approach (VanRullen, 2016). For each participant, a surrogate distribution was built by randomly shuffling the trials label and recalculating the POS for each new shuffling (number of repetitions: 1000). Then, a group-level surrogate distribution was built by randomly picking a POS sample for each participant and averaging the samples across participants (number of repetitions: 100 000). Last, a cluster-based permutation procedure was used to compare the empirical POS values to the group-level surrogate distribution (cluster threshold corresponding to the two-sided 95^th^ percentile), ensuring a correction for the multiple comparison problem.

We computed the Bayes factor for paired t-tests and Pearson correlation using the *bayesFactor* toolbox in Matlab and the *BayesFactor* package with R (Version 3.5.3), and interpreted the Bayes Factor following Jarosz & Wiley, 2014.

## Results

### Temporary and idiosyncratic biases in time perception

The participants’ task was to judge which of two stimuli in an audiovisual pair of asynchronous stimuli was presented first. In an initial screening session, we determined the idiosyncratic bias of each participant. If a participant exhibited the tendency (across a set of balanced stimulus-onset asynchronies) to perceive the flash first more often, we would deem the flash-first response (Vfirst) as matching this individual’s bias (matching-bias) and the sound-first response (Afirst) as against-bias (Fig. 1B). We computed the Point of Subjective Simultaneity (PSS) as a proxy for an individual’s bias (Grabot & Wassenhove, 2017) and the Just Noticeable Differences (JNDs) for each individual and condition (Fig. 1B). In a planned sample, 12 participants were biased towards Vfirst responses (negative PSS, mean ± SD = −44 ± 46ms) and 12 were biased towards Afirst responses (positive PSS, mean ± SD = 38 ± 31ms).

To induce a temporary bias, we tested participants’ perception after prolonged exposure to constant audio-leading or visual-leading delays of 200ms (A200V and V200A conditions). As expected, recalibration significantly changed the perceived simultaneity (Table 2). A repeated-measure one-way ANOVA showed that there was a significant effect of recalibration on PSS (F(2,23) = 3.48, p = 0.039, η_p_^2^ = 0.67; Fig. 2A). A post-hoc t-test revealed that the PSS after A200V recalibration significantly decreased compared to V200A recalibration (t(23) = −2.51, p = 0.019, CI95 = [−23, −2]ms, effect size = 12ms, BF = 2.80). In addition, the PSS significantly differed between the A200V and no recalibration conditions (t(23) = −2.34, p = 0.028, CI95 = [−29, −2]ms, effect size = 16ms, BF = 2.04), but not between the V200A and no recalibration conditions (t(23) = −0.45, p = 0.657, CI95 = [−17, 11]ms, effect size = 3ms, BF = 0.23).

**Table 2.** Group averaged PSSs and JNDs for each condition (mean ± standard deviation).

| condition | no | A200V | V200A |
| --- | --- | --- | --- |
| PSS (ms) | -3 ± 57 | -17 ± 41 | -6 ± 41 |
| JND1 (ms) | -83 ± 54 | -85 ± 57 | -70 ± 54 |
| JND2 (ms) | +77 ± 75 | +48 ± 46 | +58 ± 63 |

Recalibration also induced a change in perceptual sensitivity, as measured by the difference between JND1 and JND2 (denoted ΔJND). A repeated-measure one-way ANOVA showed that there was a significant effect of recalibration (F(2,23) = 3.85, p = 0.028). A post-hoc t-test revealed that the ΔJND during no recalibration differed significantly from the A200V (t(23) = −2.30, p = 0.031, CI95 = [−54, −3]ms, effect size = 28 ms, BF = 1.90) and V200A conditions (t(23) = −2.22, p = 0.037, CI95 = [−63, −2]ms, effect size = 32 ms, BF = 1.66), showing that the temporal order judgements became less sensitive after recalibration.

### Behavioral results from the EEG sessions

The same participants then performed the same temporal order judgment task while EEG was recorded. Here, we used only two audiovisual SOAs as stimuli per condition (no-, A200V- and V200A-conditions) defined as the individual JND1/2 extracted for each individual from the screening session. The behavioral data confirmed that the selection of JNDs for the EEG experiment was appropriate and resulted in the expected percentage of Vfirst responses (30% for JND1 and 70% for JND2; Fig. 2B): a one-sample t-test indicated that the difference between the actual percent of Vfirst responses and the *a priori* percentage was not statistically different (t(143) = 1.23, CI95 = [−0.009, 0.038], p = 0.220, BF = 0.19). Using a sound-leading (JND1) and a flash-leading (JND2) delay for each individual allowed us to orthogonalize the correctness and the biasedness of the responses, while ensuring a sufficient number of trials per response (Fig. 1D). Since our population was composed of 12 participants favoring flash-first responses and 12 participants favoring sound-first responses, the correctness and the biasedness of the responses are also orthogonal to the responses (Afirst, Vfirst).

### Alpha activity reflects processes that help to overcome an idiosyncratic bias

Given the presumed role of pre-stimulus alpha band activity in shaping subsequent perceptual responses, we quantified the relation between alpha activity and the idiosyncratic biases. To investigate whether alpha power predicted the correctness or biasedness of a subsequent response, we extracted alpha power from −600 to −200ms before the onset of the first stimulus in no recalibration condition and entered this into a 2×2 ANOVA combined with a spatio-temporal cluster-based permutation test (Fig. 3A). Importantly, the contrasts for correctness and biasedness were orthogonal given the experimental design and were controlled for the physical order of stimuli presentation. We found a significant positive cluster for biasedness over left fronto-central sensors (−598 to −438ms, cluster-value = 291.50, p = 0.044), suggesting that an increase in alpha power predicts a matching-bias response (Fig. 3A, right). No effect (even at a reduced p<0.15) was found for correctness, nor was there any significant interaction. A Bayesian analysis revealed that there is positive or substantial evidence that alpha power averaged across the left frontal cluster does not differ between correct and incorrect response (BF = 0.21), that there is no interaction effect (BF = 0.27), and confirmed decisive evidence for an effect of biasedness (BF = 112).

**Fig. 3.**
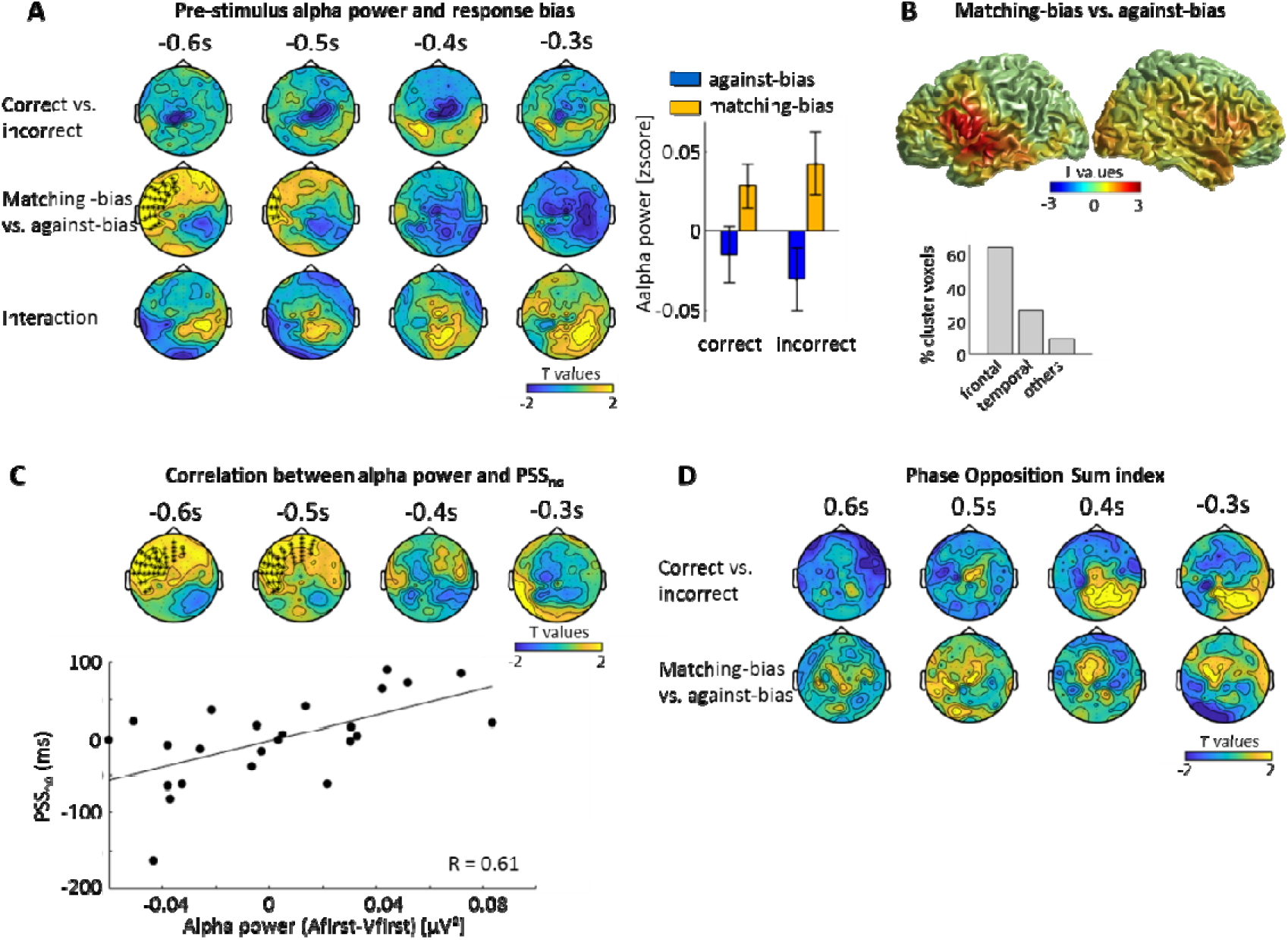
Pre-stimulus alpha power and idiosyncratic bias. **A.** Statistical maps from an ANOVA testing for main effects (correctness, matching bias) and an interaction performed for the no-recalibration condition on pre-stimulus alpha power (7.1-12.8Hz). One positive significant cluster for biasedness was found. Each topography shows the average t-map across a 100ms temporal window starting at the indicated time point. The bar graph shows alpha power within the significant cluster, for each response (mean ± SEM across participants). **B.** Source-space contrast of pre-stimulus alpha power between matching-bias and against-bias responses between −600 and −200ms. The center of mass of the cluster corresponding to uncorrected p<0.05 is located in the left rolandic operculum. 64% of the cluster covers left frontal areas, 27% left temporal areas and 9% left postcentral areas and insula (based on the AAL atlas). **C.** Correlation between PSS_no_ and the alpha difference (Afirst-Vfirst) tested using a spatio-temporal cluster based permutation test. One significant cluster was found. **D.** The phase opposition sum (POS) index was used to determine whether the instantaneous alpha phase has different concentration between two conditions. No significant cluster was found.

To localize the neural generators underlying the effect of biasedness, we performed a source reconstruction on pre-stimulus alpha activity and contrasted matching-bias and against-bias responses. This revealed a cluster in left temporo-frontal areas, consistent with the left frontal topography (Fig. 3B). The center of mass of this cluster (at an uncorrected p<0.05) was located in the left rolandic operculum (MNI coordinates: −60 +5 +6, corresponding to Brodmann area 44). Based on the AAL atlas, this cluster comprised prominent parts of the left frontal inferior operculum, rolandic operculum and temporal superior area, with 64% of the grid points contained in the cluster falling in left frontal areas, 27% in left temporal areas, and 9% in the left postcentral area and the insula (Fig. 3B).

If pre-stimulus alpha band activity indeed reflects processes that help to overcome an idiosyncratic bias, the difference in power between responses (Afirst and Vfirst) should not only show a general association with biasedness but should also scale in proportion to the strength of the individual bias, hence the magnitude of the individual PSS. To test this, we computed a Pearson correlation between the difference in pre-stimulus alpha power (Afirst-Vfirst) and the individual PSS extracted from the no recalibration condition (Fig. 3C). This revealed a significant cluster over left fronto-central sensors (−598 to −438ms, cluster-value = 436.09, p = 0.037), the location of which was consistent with the results from the ANOVA above. Within this cluster, the alpha power difference between Afirst and Vfirst response was R = 0.61 (bootstrap-based CI95% = [0.27, 0.82], R = 0.58, BF = 11, Fig. 3C). As a control, we also tested within the same cluster whether the alpha power difference between correct and incorrect response correlated with the PSS, and found no evidence for a significant correlation (R = −0.04, p = 0.864, BF = 0.16).

We explored whether this result was specific to the alpha band by repeating the above analysis for the theta, low beta and high beta bands. No significant clusters were found for biasedness in any of these bands (Fig. 4). However, high beta band activity was related to correctness (−298 to −218ms, cluster-value = −395.21, p=0.002), with decreased beta power in occipital central electrodes predicting a correct response. To investigate the amount of evidence in favor of the null hypothesis, we calculated a Bayes Factor for a paired t-test at each sensor and time point. 67.5%, 75.4% and 70.5% of the data points in the theta, low beta and high beta bands respectively showed positive and substantial evidence for the null hypothesis (BF < 1/3), while only 1.2%, 0.8% and 1.3% of the data point in the same bands showed positive and substantial evidence (BF > 3) for a difference in biasedness (Table 3). Given the large number of data points tested, we consider these few ‘significant’ data points as false positives, as their number does not exceed a threshold at α = 0.05. On the contrary, more than 5% of the data points in the high beta band showed positive and substantial evidence for a difference in correctness, in line with the significant cluster found.

**Table 3.** Bayes factors for theta and beta band results. Occurrence of (sensors, time point) couple with BF > 3, denoting a positive and substantial evidence for a difference between conditions (correct vs. incorrect or matching-bias vs. against-bias) and BF < 1/3, denoting a positive and substantial evidence for the null hypothesis (no difference).

|  | theta | low beta | high beta |
| --- | --- | --- | --- |
| **Correctness** |  |  |  |
| **BF > 3** | 2.2 % | 2.9 % | 5.4 % |
| **BF < 1/3** | 69.1 % | 60.2 % | 57.5 % |
| **Biasedness** |  |  |  |
| **BF > 3** | 1.2 % | 0.8 % | 1.3 % |
| **BF < 1/3** | 67.5 % | 75.4 % | 70.5 % |

**Fig. 4.**
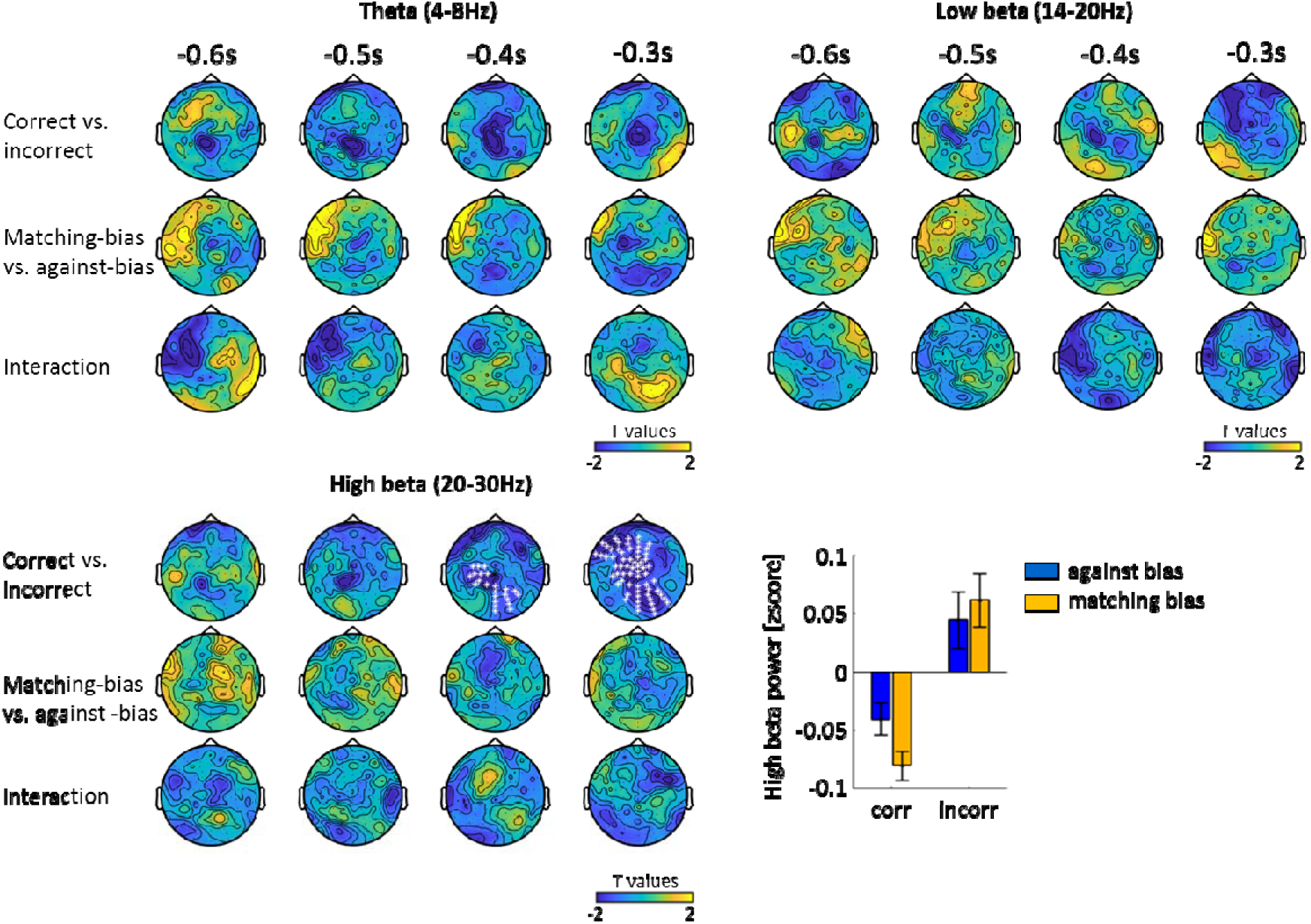
Specificity to the alpha band. We tested whether pre-stimulus power in other frequency bands (theta, low beta and high beta) was related to response correctness, biasedness and their interaction. There was no effect of biasedness in any frequency band, suggesting that this effect is specific to the alpha band. In the high 20-30Hz beta band, reduced power was predictive of a correct response (mean ± SEM across participants). See Table 3 for Bayes factors.

We also investigated whether the phase of alpha activity was predictive of whether the following response was correct or biased, since previous studies suggested a link between alpha phase, detection performance and prior expectation (Busch, Dubois, & VanRullen, 2009; Sherman, Kanai, Seth, & VanRullen, 2016). A non-parametrical permutation-based procedure on the Phase Opposition Sum (POS) index (VanRullen, 2016) revealed no significant effects for correctness or biasedness (Fig. 3D).

### Pre-stimulus alpha reflects both idiosyncratic and temporary biases

Next, we investigated the interplay between the idiosyncratic bias and the temporary biases induced by recalibration. If alpha activity is reflective of neural processes that are affected by both types of biases, we would expect that the relationship between alpha power and the biasedness of responses would remain the same across recalibration conditions. If, however, alpha indexes only an idiosyncratic bias, we would expect that the explanatory power differs between recalibration conditions, because the influence of any long-term biases is shadowed by the temporary influence of recalibration. To address this, we focused on the cluster (i.e. electrodes and time points) that showed a significant alpha effect both for the ANOVA contrast and the correlation analysis in the no recalibration condition (−598 to −438ms, 19 electrodes, Fig. 3A and 3C).

To quantify whether alpha power is predictive of either the original idiosyncratic bias, or a updated bias emerging from recalibration, we split trials from A200V and V200A conditions i) according to correctness and biasedness relative to the recalibrated biases (PSS_A200V_ and PSS_V200A_), or ii) relative to the original idiosyncratic bias (PSS_no_).We then entered these alpha power values into a linear mixed-effect model with the factors correctness, biasedness, their interaction, and participants as a random effect (Fig. 5A). When using the adapted PSSs, alpha was significantly related to biasedness (χ^2^(1) = 4.90, p = 0.027, BF = 2.06), but not to correctness (χ^2^(1) = 0.13, p = 0.715, BF = 0.22), and the interaction was not significant (χ^2^(1) = 1.06, p = 0.303, BF = 0.20). When using the non-adapted PSS, no factors were significant (biasedness: χ^2^(1) = 0.26, p = 0.611, BF = 0.24; correctness: χ^2^(1) = 0.13, p = 0.720, BF = 0.23; interaction: χ^2^(1) = 0.43, p = 0.509, BF = 0.02). Furthermore, a model comparison based on a likelihood ratio test showed that the model using the adapted PSSs (LL = 96.1) was explaining significantly more variance than the non-adapted model (LL = 94.5, χ^2^ = 2.76, p < 10^−3^). An estimated Bayes factor suggested that the model with the adapted PSSs was 10 times more likely to occur than the model with the PSS_no_. These results suggest that pre-stimulus alpha power predicts whether a subsequent response will follow or not the momentary and contextually adapted bias.

**Fig. 5.**
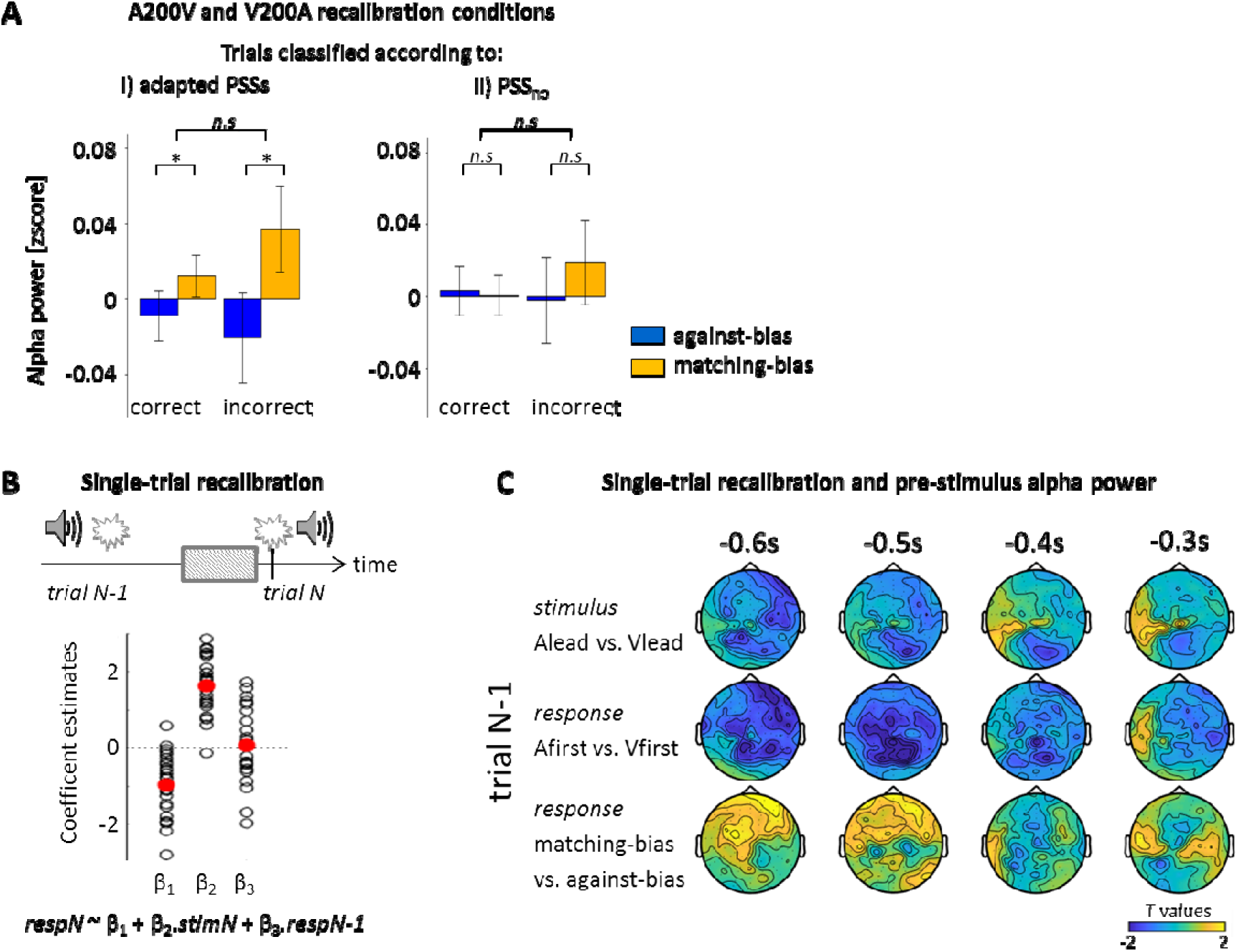
Pre-stimulus alpha power and temporary biases. We investigated whether pre-stimulus alpha power was also modulated by prolonged and short-term temporal recalibration. **A.** The trials in A200V and V200A conditions were either classified following i) the adapted PSSs (PSS_A200V_ and PSS_V200A_) or ii) the non-adapted PSS (PSS_no_). The alpha power was computed according to biasness and correctness for these two classifications and averaged across the conjunction cluster derived from the ANOVA and the correlation analysis (Fig. 3A and 3C). A two-way ANOVA (factors: biasness, correctness) showed that the biasedness factor was significant only when trials were classified according to adapted PSSs (*: p < 0.05, *n.s.*: non-significant). **B.** A single-trial model was used to explain the response in the current trial N (respN) based on the current stimulus (stimN) and the previous response (respN-1). The coefficient estimates for each individual (black dots) and the mean across participants (red) are shown. **C.** The relationship between pre-trial N alpha power and the response of trial N-1 was investigated by splitting trials according to either stimulus, response or biasedness of the response in trials N-1. No significant clusters were found.

### Trial-by-trial dependencies

Temporal recalibration is not only observed after prolonged exposure but can also emerge on a trial-by-trial basis (Van der Burg, Alais, & Cass, 2013; Van der Burg, Alais, & Cass, 2015). We investigated whether and how the previous trial influenced the alpha band activity prior to the subsequent trial and contributed to the behavioral response in the subsequent trial. First, we asked whether the previous stimulus (stimN-1) or the previous response (respN-1) significantly influenced the subsequent response (respN), by comparing the ability of distinct linear models based on different combinations of these predictors to predict the subsequent response (Table 1). The model providing the most parsimonious account of the data (according to the group-level BIC criterion) was the model with stimN and respN-1 as predictors (BIC = 43 848, Table 1). The model estimates and t-values averaged across participants were β(intercept) = −0.97 ± 0.84, p < 0.05 for 21/24 participants; β(stimN = Alead) = 1.63 ± 0.76, p < 0.05 for 23/24 participants; β(respN-1 = Afirst) = 0.06 ± 0.97, p<0.05 for 20/24 participants; df = 1 546 ± 70 (mean ± SD, Fig. 5B). Because a previous study has suggested that trial-by-trial temporal recalibration is driven by the previous stimulus rather than the previous response (Van der Burg, Alais, & Cass, 2018), we implemented a second analysis to better contrast the influence of stimN-1 and respN-1. For this, we removed the natural correlation between these two predictors by equalizing, for each participant, the number of trials between responses (Afirst or Vfirst) for a given previous stimulus (Alead or Vlead), and recomputed the linear model with these predictors. Still, stimN-1 was a significant predictor for only 6 out of 24 participants (β= −0.143 ± 0.285, t = −0.801 ±1.529, p<0.05 for 6 participants), while respN-1 was significant for most participants (β= 0.157 ± 0.962, t=1.062 ± 5.945, p<0.05 for 21 participants; df = 947 ± 203; mean ± SD). Hence in the present data, subsequent responses are more tied to the previous response than the previous stimulus.

We then asked whether the alpha activity before the subsequent trial was related to the stimuli or behavior in the previous trial (Fig. 5C). We sorted trials according to either the stimulus order (Alead vs. Vlead), the response (Afirst vs. Vfirst), and the biasedness (matching-bias vs. against-bias) of the previous trial. A spatio-temporal cluster-based permutation test revealed no significant effects. In particular, for biasedness, one cluster over frontal electrodes did not pass the significance threshold at 0.05 (p = 0.111; −498 to −418ms; Bayesian analysis: 57.5% of the data points with BF < 1/3, 0.7% with BF > 3), suggesting that the previous stimuli or responses did not influence the pre-stimulus alpha activity for the subsequent trial.

Using additional analyses we further ruled out that alpha mediates, or modulates, the influence of the response in the previous trial on the current trial: A mediation analysis on alpha power from the left frontal cluster, revealed no significant mediation (path from respN-1 to the alpha power: a=0.022, p = 0.051; path from the alpha power to the respN: b=0.05, p =0.031; mediation term: ab=5.2e-5, p=0.424). Further, logistic modelling of the subsequent response revealed no interaction between alpha and the previous response (β(intercept) = - 0.01 ± 0.53, p < 0.05 for 20/24 subj; β(respN-1=Alead) = 0.07 ± 0.78, p < 0.05 for 20/24 subj; β(alpha) = 0.04 ± 0.07, p < 0.05 for 2/24 subj, β(alpha*respN-1) = −0.04 ± 0.12, p < 0.05 for 3/24 subj; df= 1 546 ± 70; mean ± SD).

## Discussion

Idiosyncratic biases are ubiquitous in human behavior but are often ignored in experimental work (Matthews & Meck, 2014; Kosovicheva & Whitney, 2017; Ipser, et al., 2017). Their origin remains unknown but the persistency of these biases over time suggests that they arise from structural or functional characteristics of an individual’s brain (Kanai & Rees, 2011). Indeed, previous work has speculated about the relation of such biases with inter-individual differences in genotypes, neurotransmitters levels, brain structure or signatures of spontaneous brain activity (Kanai & Rees, 2011; Mennes, et al., 2011; Kleinschmidt, Sterzer, & Rees, 2012; Romei, Murray, & Thut, 2013; Matthews & Meck, 2014; Haegens, Cousjin, Wallis, Harrison, & Nobre, 2014; Marshall, Bergmann, & Jensen, 2015; Chechlacz, Gillebert, Vangkilde, Petersen, & Humphreys, 2015).

We here focus on a prevailing signature of spontaneous brain activity, alpha band oscillations, and tested the previously speculated hypothesis that alpha band activity relates to the degree of how biased an individual response is (Grabot, Kösem, Azizi, & Wassenhove, 2017). Our results show that a decrease in frontal pre-stimulus alpha power is associated with an increased chance that the subsequent response will go against an idiosyncratic bias, but is not predictive of whether the response will be correct. Further, the stronger an individual idiosyncratic bias, the more alpha power will differentiate between responses that match and mismatch the bias. Importantly, this relation also holds after temporal recalibration, a manipulation that affects contextual short-term biases in perception. Finally, we found that this relation is specific to the alpha band. Hence, our results show that frontal pre-stimulus alpha band activity indexes processes that help to overcome an individual’s momentary, rather than only a stable and long-term, bias.

### A mechanistic role of alpha power

Previous studies have dissociated two patterns of alpha activity typically described in relation to perception: a frontal alpha involved in cognitive control, and a parieto-occipital alpha involved in spatial perception (Noonan et al., 2016; Sadaghiani et al., 2016; Wöstmann et al., 2019). We suggest that the alpha activity observed here is part of a frontal network and discuss its function in relation to perceptual decisions within this framework.

Parieto-occipital alpha is usually observed in studies on spatial attention, which show that lateralized changes in alpha power are predictive of subsequent target detection performance (Sauseng et al., 2005; Thut et al., 2006; Romei et al., 2010). Recent studies tried to define the point of action of alpha activity in the framework of signal detection theory (Iemi et al., 2017; Iemi & Busch, 2018). These suggest that alpha reflects an additive mechanism affecting sensory encoding more than the decision process (Limbach & Corbalis, 2016; Iemi et al., 2017), and relates more to subjective awareness and confidence than perceptual accuracy (Samaha et al., 2017, Wöstmann et al., 2018; Iemi & Busch, 2018). Given that alpha power correlates with sensory neural firing rates (Jensen & Mazaheri, 2010; Klimesch, Sauseng, & Hanslmayr, 2007; Haegens et al., 2011; Klimesch, 2012), alpha power can be interpreted as reflecting the overall excitability in those regions encoding the sensory information. Our experiment was designed to dissociate a response bias from performance and does not allow an unambiguous interpretation in the framework of signal detection theory. However, the prominent localization of alpha effects over frontal sites found here is rather suggestive of an origin that is distinct from this parietal alpha.

Frontal alpha has been linked to cognitive control and the selection of task-relevant information (Sauseng, et al., 2005; Sadaghiani & Kleinschmidt, 2016; Wöstmann, Alavash, & Obleser, 2019). Thereby, frontal alpha may affect decision-making independently of the sensory evidence, for example, by adding a fixed proportional probability to choose one response over the other. In that case, the response bias could change without changing the overall fraction of correct responses. Still, at a mechanistic level this frontal origin and its functional interpretation can be linked to an understanding of the possibly multiple roles of alpha band activity in perception: in a complete sensory-decision cascade, parietal alpha may affect early sensory encoding (Lou, Li, Philiastides, & Sajda, 2014; Iemi, Chaumon, Crouzet, & Busch, 2017), while frontal alpha may affect the selection of a specific response independently of the outcome of the preceding sensory computation.

In terms of the underlying neurobiological correlates, idiosyncratic biases have been suggested to arise from individual variability in specific structural brain features or functional connectivity (Freeman, et al., 2013; Grabot & Wassenhove, 2017; Ipser, Karlinski, & Freeman, 2018). These structuro-functional differences may have been shaped by developmental experience, hence introducing individual differences in the sensory decision-making process. Similarly, cognitive control rests on consistent rules learned from past experience (Miller, 2000). One could therefore speculate that changes in frontal alpha reflect the degree to which such structuro-functional imbalances shape behavior and are reflective of how cognitive control impacts perceptual reports. How precisely frontal alpha band activity affects the degree of biasedness of perceptual reports remains to be investigated, similar as the direct links between alpha band activity, structuro-functional connectivity and how these relate to cognitive control.

### Decreased beta power predicts correct responses

We found that decreased beta power (20-30Hz) was predictive of correct responses. This fits with results obtained using similar audiovisual simultaneity judgment tasks and with several studies relating beta decrease to a better sensory encoding (Griffiths et al., 2019) and beta increase to illusory (hence incorrect) perception (see Keil et al, 2018, for a review; Kaiser et al., 2019). However, one study reported an opposite relationship between accuracy and beta power in a spatial auditory task (Bernasconi et al., 2011), calling for more systematic studies on how the nature of stimuli and task influence the neural generators of relevant beta activity and its influence on performance.

### The apparent asymmetry of temporal recalibration

We found no significant effect of VA recalibration on participants’ PSS. This is in contrast to AV recalibration, which significantly reduced the PSS. Such an asymmetry between AV and VA recalibration has been observed consistently in multiple studies (Fujisaki, Shimojo, Kashino, & Nishida, 2004; Vroomen, Keetels, Gelder, & Bertelson, 2004; Kösem, Gramfort, & Wassenhove, 2014; Ikumi & Soto-Faraco, 2014), yet a clear explanation remains missing. One possibility is that in everyday life we are mainly confronted with VA asynchronies, as light travels faster than sound and for many actions the visual movement precedes any produced sound. The consistency of this observation across studies suggests that our brain may indeed be differentially sensitive to AV and VA adaptation, in the same way that integrating audiovisual events leads to an asymmetric temporal binding window (van Wassenhove, Grant, & Poeppel, 2007; Stevenson, Zemtsov, & Wallace, 2012).

### Distinct mechanisms mediating prolonged and single-trial recalibration

The neural correlates of temporal recalibration are still poorly understood (Stekelenburg, Sugano, & Vroomen, 2011; Kösem, Gramfort, & Wassenhove, 2014; Simon, Noel, & Wallace, 2017; Simon, Nidiffer, & Wallace, 2018). We found that pre-stimulus alpha activity predicts the degree of biasedness to perception after prolonged temporal recalibration, but does not mediate trial-by-trial recalibration, i.e. the influence of a previous response on behavior (Van der Burg, Alais, & Cass, 2013). This suggests that distinct mechanisms underlie prolonged and single-trial recalibration, a conclusion supported by psychophysical experiments in the temporal (Van der Burg, Alais, & Cass, 2015) and the spatial domains (Bruns & Röder, 2015; Bruns & Röder, 2017; Watson, Akeroyd, Roach, & Webb, 2019). A role of alpha activity in reflecting an individual’s overall perceptual tendency without interacting with trial-by-trial dependencies is also supported by a recent study on multisensory integration (Rohe, Ehlis, & Noppeney, 2019). Assuming that alpha reflects the influence of cognitive control, the dissociation of alpha activity and trial-by-trial recalibration could reflect the need to strike a balance between the sustainability of a temporally consistent internal state, and the flexibility required to cope with dynamic environments. Frontal alpha activity would help sustaining internal states in line with long-term experience without interfering with other trial-by-trial adaptive mechanisms that also shape perception.

Alternatively, the perceptual dependencies often described as single-trial recalibration may in fact not be a genuine recalibration effect but reflect a bias to repeat a previous response in consecutive trials (Roseboom, 2019; Keane, Bland, Matthews, Carroll, & Wallis, 2019). Our data revealed a significant choice-repetition bias, which is inconsistent with a recalibration effect (exposing participants to audio-leading stimuli would increase their tendency for Vfirst responses). However, since a genuine recalibration effect can be observed when one controls for choice-repetition biases (Keane, Bland, Matthews, Carroll, & Wallis, 2019), further work is needed to determine the neural mechanisms underlying single-trial temporal recalibration. In the spatial domain one study recently suggested that recalibration induced by discrepant multisensory information arises from medial parietal brain regions involved in episodic and spatial memory (Park & Kayser, 2019), suggesting that in the spatial domain single-trial and long-term recalibration emerge from distinct neural structures (Bruns & Röder, 2010; Bruns, Spence, & Roeder, 2011; Bosen, Fleming, Allen, O’Neill, & Paige, 2017). Still, further studies are required to directly compare the neural mechanisms underlying different time scales of adaptive behavior within and across different domains of sensory information.

## Acknowledgments

This work was supported by the European Research Council (to CK ERC-2014-CoG; grant n°646657).

